# Immune Evasion, Cell-Cell Fusion, and Spike Stability of the SARS-CoV-2 XEC Variant: Role of Glycosylation Mutations at the N-terminal Domain

**DOI:** 10.1101/2024.11.12.623078

**Authors:** Pei Li, Julia N. Faraone, Cheng Chih Hsu, Michelle Chamblee, Yajie Liu, Yi-Min Zheng, Yan Xu, Claire Carlin, Jeffrey C. Horowitz, Rama K. Mallampalli, Linda J. Saif, Eugene M. Oltz, Daniel Jones, Jianrong Li, Richard J. Gumina, Joseph S. Bednash, Kai Xu, Shan-Lu Liu

**Affiliations:** Center for Retrovirus Research, The Ohio State University, Columbus, OH 43210, USA; Department of Veterinary Biosciences, The Ohio State University, Columbus, OH 43210, USA; Molecular, Cellular, and Developmental Biology Program, The Ohio State University, Columbus, OH 43210, USA; Texas Therapeutic Institute, Institute of Molecular Medicine, University of Texas Health Science Center at Houston, Houston, TX 77030, USA; Department of Internal Medicine, Division of Cardiovascular Medicine, The Ohio State University, Columbus, OH 43210, USA; Department of Internal Medicine, Division of Pulmonary, Critical Care, and Sleep Medicine, The Ohio State University, Columbus, OH 43210, USA; Dorothy M. Davis Heart and Lung Research Institute, The Ohio State University, Wexner Medical Center, Columbus, OH 43210, USA; Center for Food Animal Health, Animal Sciences Department, OARDC, College of Food, Agricultural and Environmental Sciences, The Ohio State University, Wooster, OH 44691, USA; Veterinary Preventive Medicine Department, College of Veterinary Medicine, The Ohio State University, Wooster, OH 44691, USA; Viruses and Emerging Pathogens Program, Infectious Diseases Institute, The Ohio State University, Columbus, OH 43210, USA; Department of Microbial Infection and Immunity, The Ohio State University, Columbus, OH 43210, USA; Pelotonia Institute for Immuno-Oncology, The Ohio State University Comprehensive Cancer Center Arthur G James Cancer Hospital and Richard J Solove Research Institute, Columbus, Ohio, USA; Department of Pathology, The Ohio State University Wexner Medical Center, Columbus, OH, USA; Department of Physiology and Cell Biology, College of Medicine, The Ohio State University Wexner Medical Center, Columbus, OH 43210, USA

## Abstract

SARS-CoV-2 continues to evolve, producing new variants that drive global COVID-19 surges. XEC, a recombinant of KS.1.1 and KP.3.3, contains T22N and F59S mutations in the spike protein’s N-terminal domain (NTD). The T22N mutation, similar to the DelS31 mutation in KP.3.1.1, introduces a potential N-linked glycosylation site in XEC. In this study, we examined the neutralizing antibody (nAb) response and mutation effects in sera from bivalent-vaccinated healthcare workers, BA.2.86/JN.1 wave-infected patients, and XBB.1.5 monovalent-vaccinated hamsters, assessing responses to XEC alongside D614G, JN.1, KP.3, and KP.3.1.1. XEC demonstrated significantly reduced neutralization titers across all cohorts, largely due to the F59S mutation. Notably, removal of glycosylation sites in XEC and KP.3.1.1 substantially restored nAb titers. Antigenic cartography analysis revealed XEC to be more antigenically distinct from its common ancestral BA.2.86/JN.1 compared to KP.3.1.1, with the F59S mutation as a determining factor. Similar to KP.3.1.1, XEC showed reduced cell-cell fusion relative to its parental KP.3, a change attributed to the T22N glycosylation. We also observed reduced S1 shedding for XEC and KP.3.1.1, which was reversed by ablation of T22N and DelS31 glycosylation mutations, respectively. Molecular modeling suggests that T22N and F59S mutations of XEC alters hydrophobic interactions with adjacent spike protein residues, impacting both conformational stability and neutralization. Overall, our findings underscore the pivotal role of NTD mutations in shaping SARS-CoV-2 spike biology and immune escape mechanisms.

## INTRODUCTION

Despite the fact that the COVID-19 pandemic appears to be moving into a more endemic phase (1–3), SARS-CoV-2 continues to mutate and generate new variants with corresponding waves of infection (4). The BA.2.86 lineage of SARS-CoV-2 emerged in 2023 and marked a new evolutionary turning point for the virus due to its possession of over 30 mutations distinct from the previously dominant XBB.1.5 (4–11). Since then, the BA.2.86-derived JN.1 variant, defined by the additional L455S spike mutation, has largely dominated worldwide (8, 10, 12) (**Fig. 1A-C**). During the summer of 2024, this variant has been supplanted by the KP.3.1.1 variant, which is defined by the additional spike mutations F456L, Q493E, and V1104L (KP.3) and DelS31(13–15). We have recently demonstrated KP.3.1.1 spike is characterized by marked evasion of vaccinated and convalescent sera, as well as exhibiting decreased fusogenicity in CaLu-3 cells (14). These effects are largely driven by the single DelS31 mutation at the N-terminal domain (NTD) of the spike, which generates a new glycosylation site (NFT) at N30 and, in turn, dictates antigenicity and increases the stability of spike by causing the RBD to favor the down position (14, 15).

**Figure 1:**
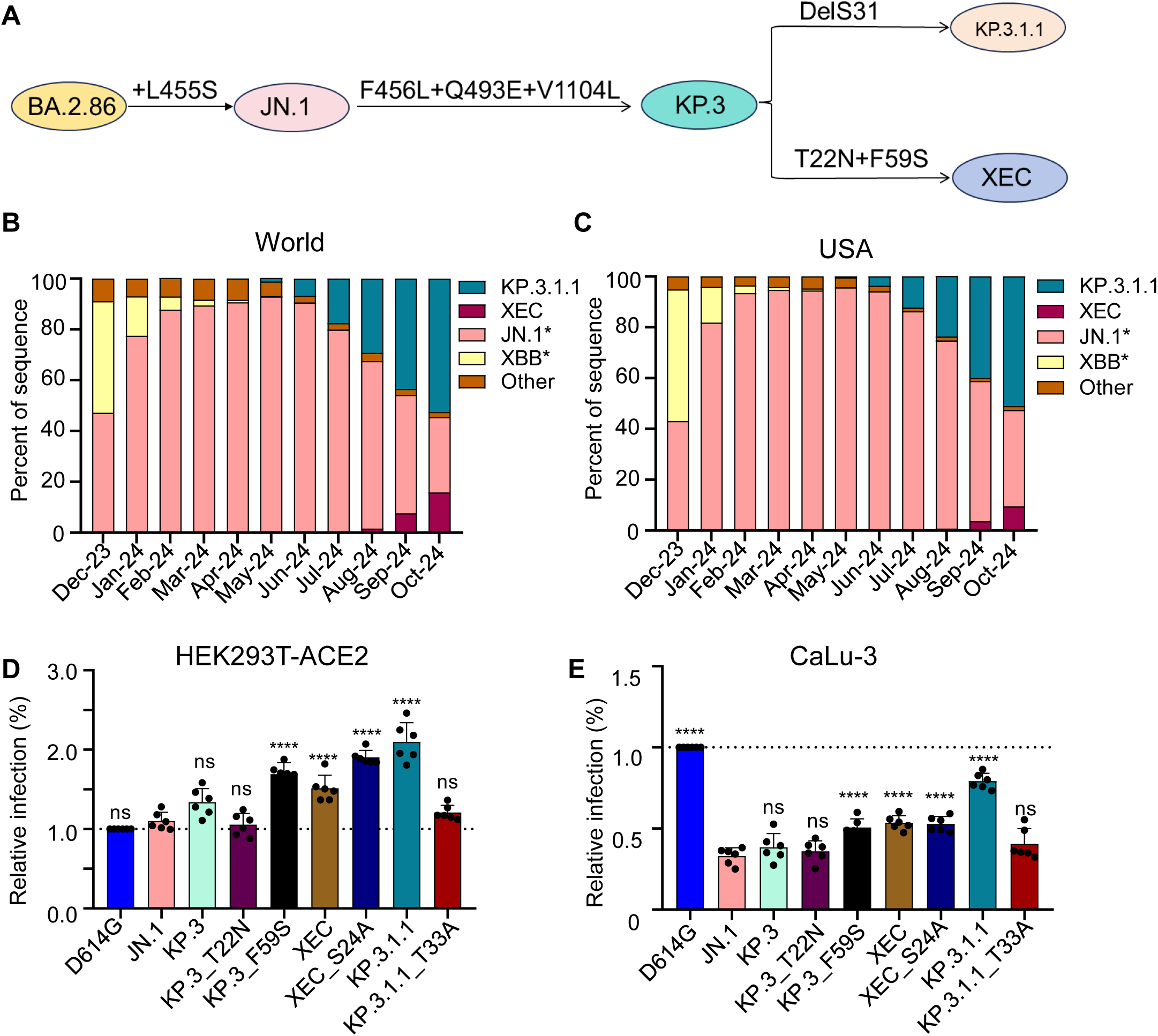
Mutations, circulation, and infectivity for JN.1-lineage variants XEC, KP.3.1.1 and KP.3. **(A)** Schematic depiction of spike-defining mutations and relationships of JN.1, KP.3, KP.3.1.1, and XEC. **(B-C)** Frequency of sequences of KP.3.1.1, XEC, JN.1, and XBB worldwide **(B)** and the United States **(C)** represented by percentage. **(D-E)** Relative infectivity of lentivirus pseudotypes bearing spikes of interest as determined by secreted *Gaussia* luciferase, with D614G set to 1.0 for comparison, in HEK293T cells expressing human ACE2 **(D)** and CaLu-3 cells **(E)**. Bars represent means with standard deviation of 3 biological replicate and 6 separate luciferase readings. Significance was determined by repeated measures one-way ANOVA in comparison to JN.1 and represented as ns p > 0.05 and ****p < 0.0001.

A new variant, XEC, is now beginning to rise in circulation. XEC is thought to be a recombinant variant between the KP.3-lineage variants K.S.1.1 and KP.3.3. The proposed split point for the recombination is in the NTD of spike, bringing together the unique NTD mutations of these two variants (16). Relative to KP.3, the XEC variant possesses the T22N and F59S mutations in spike (**Fig. 1A-C**). Like the DelS31 in KP.3.1.1, the T22N mutation in XEC is predicted to create a new glycosylation site (17). Given the dramatic effects of the single NTD mutation DelS31 in KP.3.1.1, along with those of related variants KP.2.3 and LB.1 (14), it is critical to characterize these new single mutations, especially the potential role of glycosylation in spike biology and neutralization escape.

In this study, we seek to understand the spike biology of XEC by investigating its infectivity in HEK293T-ACE2 and CaLu-3 cells, its neutralization by sera from bivalent vaccinated healthcare workers (HCWs), BA.2.86/JN.1-wave infected individuals, and XBB.1.5-vaccinated hamsters. We also characterized its fusogenicity in HEK293T-ACE2 and CaLu-3 cells, its surface expression, its furin processing, as well as its stability via S1 shedding experiments. These aspects of spike biology are evaluated alongside those of parental variants including D614G, JN.1, and KP.3, as well as the currently dominating KP.3.1.1. To enhance this analysis, we investigated the role of the single mutations that define XEC (e.g. KP.3_T22N and KP.3_F59S), especially the role of new glycosylation sites through mutations that ablate the corresponding sites in XEC (XEC_S24A), alongside a similar KP.3.1.1 mutation (KP.3.1.1_T33A). To gain structural insights, we conducted homology modeling to help better understand the residues that may play a role in spike stability and neutralization. Overall, our findings define a critical role for F59S in dictating virus infectivity and neutralization escape, as well as a role of T22N in dictating fusion and antigenicity.

## RESULTS

### F59S drives the increased infectivity of XEC

We assessed the infectivity and entry of lentiviral pseudotypes bearing each of the spikes of interest in HEK293T cells overexpressing human ACE2 (293T-ACE2) (**Fig. 1D**), and in human lung epithelial cell line CaLu-3 (**Fig. 1E**). In HEK293T-ACE2 cells, XEC exhibited a relatively modest increase in infectivity compared to JN.1 (1.4-fold, p < 0.0001) and KP.3 (1.1-fold, p = 0.11). Consistent with our previous results(14), KP.3.1.1 exhibited significantly increased infectivity relative to both JN.1 (1.9-fold, p < 0.0001) and KP.3 (1.6-fold, p < 0.0001) (**Fig. 1D**). We also assessed the impact of the single mutations, i.e., T22N and F59S, that influence the infectivity of XEC relative to KP.3. We found that the infectivity of KP.3_T22N was comparable to KP.3 (p > 0.05) while KP.3_F59S exhibited a 1.3-fold increase (p < 0.01) relative to KP.3 (**Fig. 1D**). Like KP.3.1.1 (relative to KP.3) (14, 17), KP.3_T22N introduces a potential N-linked glycosylation site into the XEC spike. To test the role of these new glycosylation sites, we generated corresponding mutations that ablated these potential glycosylation sites for each variant, e.g. XEC_S24A and KP.3.1.1_T33A, respectively. The infectivity of XEC_S24A was higher than XEC, with a 1.3-fold (p < 0.001) increase. In contrast, the infectivity of KP.3.1.1_T33A was lower than KP.3.1.1 (0.6-fold, p < 0.0001), falling to be comparable with JN.1 (p > 0.05) (**Fig. 1D**).

In CaLu-3 cells, all Omicron lineage variants had significantly lower infectivity than D614G, consistent with our previous results (**Fig. 1E**) (6, 14, 18, 19). Compared to parental KP.3, XEC exhibited a modestly increased infectivity of 1.4-fold (p < 0.01) (**Fig. 1E**). Similar to HEK293T-ACE2 cells, this increase was driven by F59S which showed an increase of 1.3-fold (p < 0.05) relative to KP.3, while T22N remained comparable to JN.1 (p > 0.05) (**Fig. 1E**). As we have shown previously (6, 20, 21), KP.3.1.1 exhibited a marked increase, about 2-fold (p < 0.0001), in infectivity relative to KP.3 (**Fig. 1E**). For glycosylation mutants, XEC_S24A remained comparable to its parental XEC, whereas the KP.3.1.1_T33A was notably lower than KP.3.1.1, which was similar to results in HEK293T-ACE2 cells (**Fig. 1E**). Overall, the F59S mutation appears to play an important role in increasing the infectivity of XEC compared to KP.3, while the N-linked glycosylation mutation DelS31 in KP.3.1.1, but not that of a similar mutation T22N in XEC, contributes to the increased infectivity in both HEK293T-ACE2 and CaLu-3 cells.

### XEC exhibits strong neutralization escape by bivalent vaccinated sera

Next, we investigated nAb titers in different cohorts, the first of which were individuals that received 3 doses of monovalent mRNA vaccine plus 1 dose of bivalent (WT+BA.4/5) mRNA vaccine (n=8) (**Fig. 2A-B)**. As we have shown previously (11, 14), JN.1, KP.3, and KP.3.1.1 all display dramatic reductions in neutralization relative to D614G. XEC also exhibited a marked decrease in nAb titers of 3.8-fold (p < 0.05) relative to JN.1, which was comparable to KP.3.1.1 (**Fig. 2A-B)**. This decrease is driven by F59S, which exhibited a 3.0-fold drop (p < 0.05) relative to JN.1, while T22N remained comparable to JN.1 (p > 0.05). Decreased neutralization titers of XEC were modestly recovered upon ablation of the acquired glycosylation site, with XEC_S24A showing a 1.2-fold increase relative to XEC (p = 0.72) (**Fig. 2A-B)**. The decreased neutralization of KP.3.1.1 was more markedly recovered with removal of the potential glycosylation site, with KP.3.1.1_T33A exhibiting a 2.0-fold increase (p = 0.13) relative to KP.3.1.1 (**Fig. 2A-B**).

**Figure 2:**
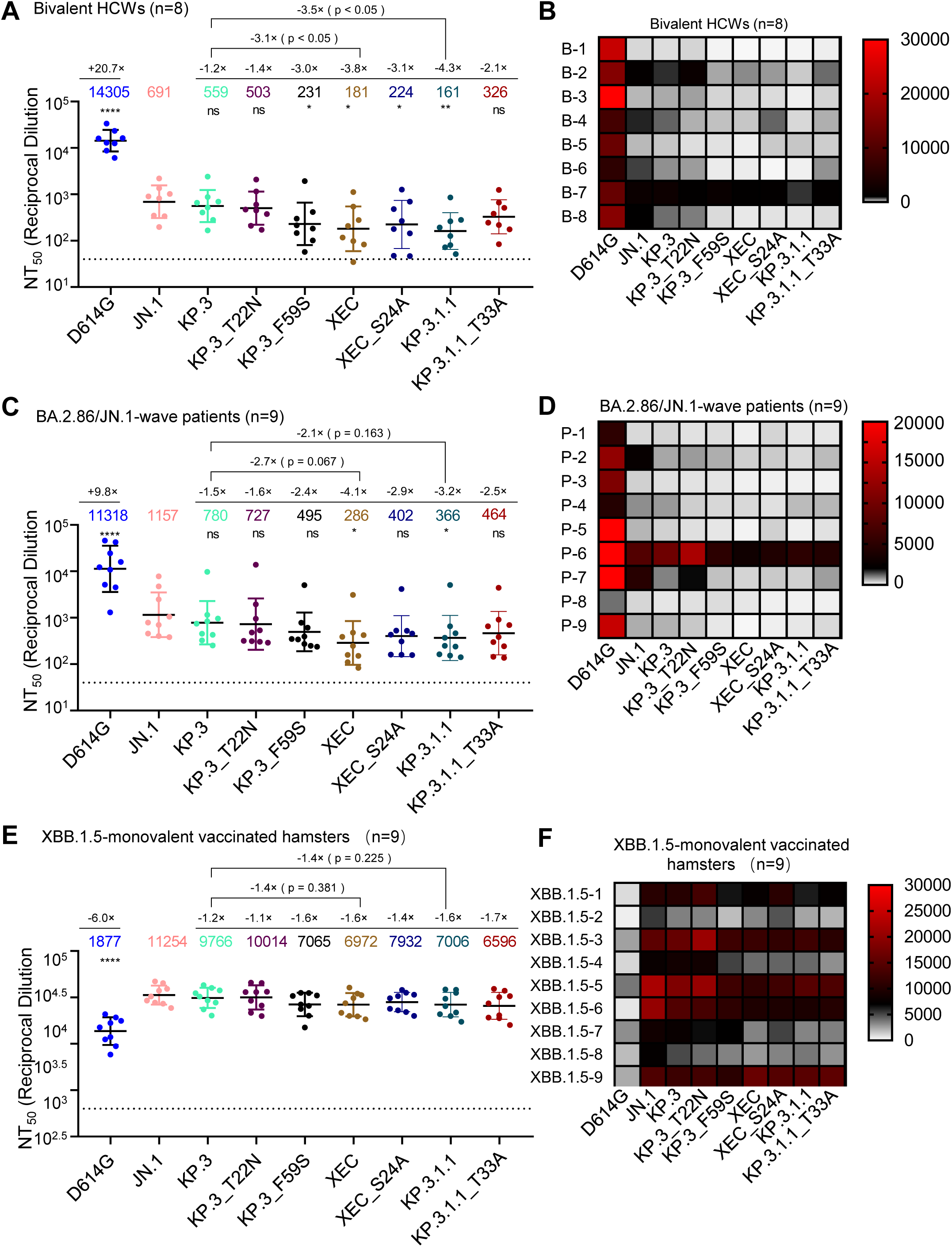
XEC exhibits strong nAb escape. NAb titers were determined using a HIV-1 pseudotyped vector neutralization assay for three cohorts of sera. **(A-B)** OSU Wexner Medical Center healthcare workers that received 3 doses of monovalent (WT) mRNA vaccine and 1 dose of bivalent (WT + BA.4/5) mRNA vaccine (n=8). **(C-D)** COVID-19 patients at the OSU Wexner Medical Center that were admitted during the BA.2.86/JN.1 wave of infection in Columbus, OH (n=9). **(E-F)** Golden Syrian hamsters vaccinated twice with a monovalent, recombinant Mumps XBB.1.5 mRNA vaccine (n=9). **(A, C, and E)** Plots represent geometric mean nAb titers at 50% (NT_50_) with standard error. Geometric mean values are listed at the top of the plots and significance was determined in comparison to JN.1 unless otherwise noted. Fold changes relative to JN.1 are listed above the geometric mean values. **(B, D, and F)** Heatmaps depicting the nAb titers for each individual in each cohort. Significance was determined in **(A, C, and E)** using log10 transformed NT_50_ values using repeated measures one-way ANOVA and represented as ns p > 0.05, *p < 0.05, and ****p < 0.0001.

### XEC exhibits the lowest neutralization for sera from BA.2.86/JN.1 infected individuals

The next cohort we investigated included sera from people infected during the BA.2.86/JN.1 wave in Columbus, Ohio, USA (n=9) (**Fig. 2C-D**). As seen previously (11, 14, 21), all variants demonstrated significantly reduced neutralization titers relative to D614G and JN.1. XEC exhibited the largest decrease, with a 4.0-fold drop in neutralization titer compared to JN.1 (p < 0.05). KP.3.1.1 showed a 3.2-fold drop relative to JN.1 (p < 0.05) (**Fig. 2C-D**). The decrease in XEC was largely driven by the F59S mutation, which exhibited a 2.3-fold decrease relative to JN.1 (p > 0.05); the T22N mutation contributed to a more modest decrease of 1.6-fold (p > 0.05) (**Fig. 2C-D**). As seen in the bivalent mRNA vaccinee cohort, titers against XEC were modestly recovered upon ablation of the new glycosylation site, with XEC_S24A exhibiting a 1.4-fold increase from XEC (p = 0.50) and 2.9-fold drop in titer relative to JN.1 (p > 0.05) (**Fig. 2C-D**). A similar effect was seen for KP.3.1.1, with KP.3.1.1_T33A exhibiting a 1.3-fold increase from KP.3.1.1 (p = 0.65) and a 2.5-fold drop in titer relative to JN.1 (p > 0.05) (**Fig. 2C-D**). Overall, these results suggest that the glycosylation mutations in the NTD of XEC and KP.3.1.1 spike contribute to the nAb evasion.

### XBB.1.5-vaccinated hamster sera robustly neutralizes all JN.1 subvariants and is modestly reduced for XEC

The final sera tested were from golden Syrian hamsters vaccinated with two doses of a recombinant mumps virus vaccine expressing XBB.1.5 spike (n=9) (**Fig. 2E-F**). As shown previously (11, 14, 21), these sera robustly neutralized all JN.1-lineage variants with much higher titers than D614G. XEC exhibited a modest decrease in titer of 1.6-fold relative to JN.1 (p > 0.05). This decrease was again driven primarily by F59S, which exhibited a 1.6-fold decrease relative to JN.1 (p > 0.05), while T22N was comparable to KP.3 and JN.1 (**Fig. 2E-F**). Ablation of the glycosylation site in XEC afforded better neutralization, with XEC_S24A exhibiting a 1.1-fold increase relative to XEC (p > 0.05). This trend was different for KP.3.1.1, with its ablating mutation KP.3.1.1_T33A actually showing a slight 1.1-fold drop relative to KP.3.1.1 (p > 0.05). Overall, titers in this cohort were much more comparable among JN.1 variants.

### Glycosylation at the NTD of XEC and KP.3.1.1 spikes differentially impacts antigenicity

To further analyze our neutralization results, we conducted antigenic cartography analysis (**Fig. 3**). Briefly, this method takes raw neutralization titer outputs and performs principal component analysis to plot titers by antigen (circles) and sera (square) in two-dimensional space in order to visualize overall trends in antigenicity where 1 antigenic distance unit (AU) is equivalent to about a 2-fold change in neutralization titer (22, 23). The panels were divided by each cohort used for the neutralization assay. In the bivalent vaccinated cohort (**Fig. 3A**), XEC displayed the farthest antigenic distance from D614G, with a distance of 7.5 AU compared to JN.1 and KP.3.1.1, which had a distance of 4.9 AU and 7.3 AU to D614G, respectively. Two single mutations of XEC, i.e., KP.3_T22N and KP.3_F59S, exhibited distinct distances compared to JN.1, with KP.3_T22N much closer (1.3 AU) whereas KP.3_F59S was more distant (3.6 AU) (**Fig. 3A)**, consistent with the more critical role of the latter in neutralization escape (**Fig. 2A-B**). Notably, the glycosylation mutants for XEC and KP.3.1.1 clustered distinctly from their parental variants, with KP.3.1.1_T33A closer to JN.1 (1.2 AU vs. 5.0 AU for KP.3.1.1) and XEC_S24A further away from JN.1 (4.8 AU vs. 3.3 AU for XEC) (**Fig. 3A, Fig. S1A**).

**Figure 3:**
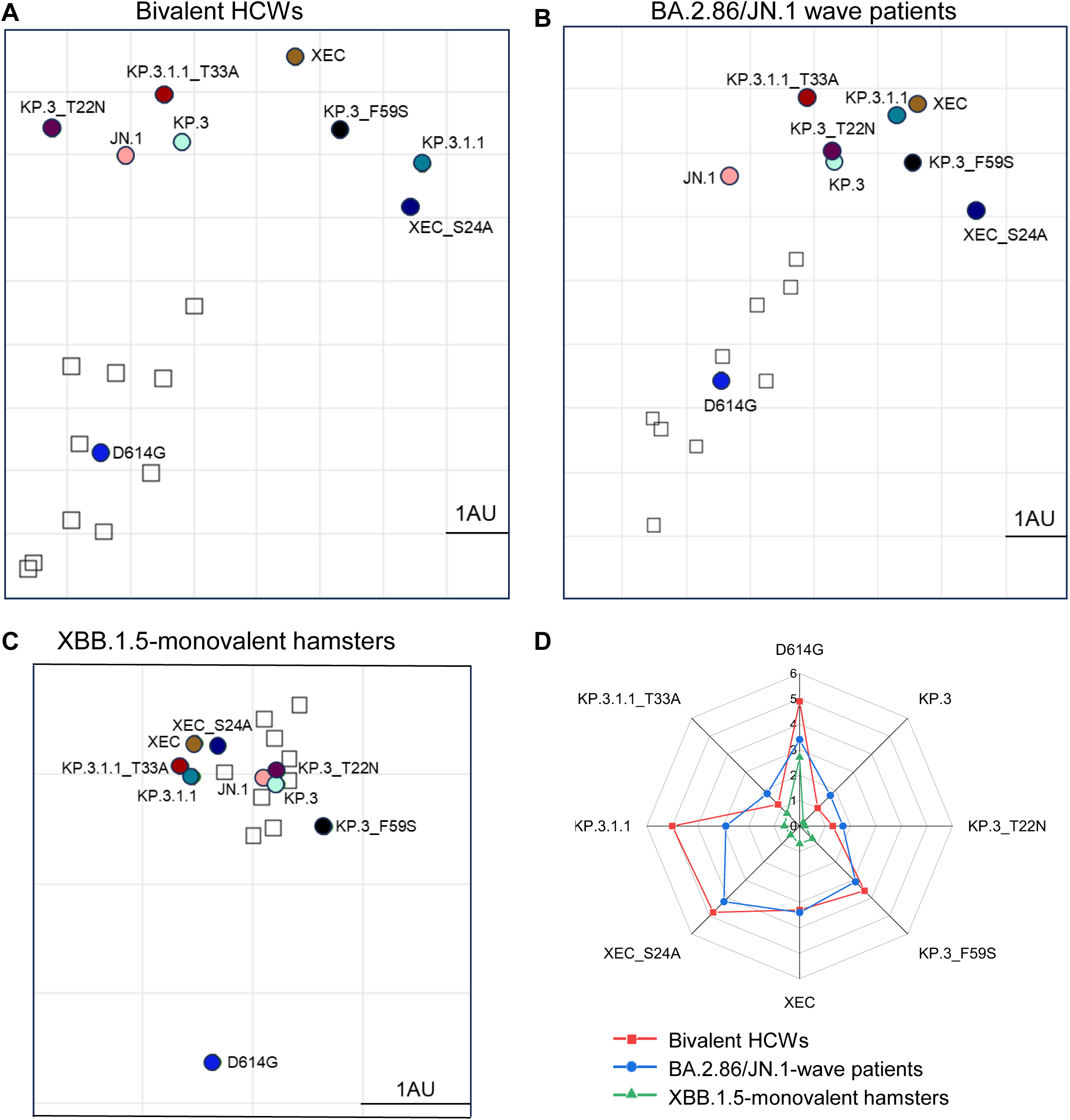
Analysis of antigenicity of XEC and related variants. **(A-C)** Antigenic cartography analysis was conducted for the nAb titers results for the bivalent vaccinated HCWs **(A)**, the BA.2.86/JN.1 wave infected patients **(B)**, and the XBB.1.5-monovalent vaccinated hamsters **(C)**. One antigenic distance unit (AU) (AU = 1) represents an approximate two-fold change in NT_50_. Circles represent the different spike antigens while boxes represent individual sera samples. **(D)** The antigenic distances of each variant relative to JN.1 from three groups of cohorts (n=3) were averaged and plotted. The scale bar represents 1 antigenic distance unit (AU).

The BA.2.86/JN.1 infected cohort displayed an overall shorter distance for JN.1 variants from D614G, with an average distance of 4.7 AU, compared to the bivalent cohort (**Fig. 3B, Fig. S1B**). Notably, XEC exhibited the largest distance from D614G and JN.1, with a distance of 5.7 AU and 3.4 AU, respectively. The single mutations KP.3_T22N and KP.3_F59S were slightly closer to D614G, with 4.3 AU and 4.9 AU, respectively. Interestingly, KP.3_T22N was much closer to parental KP.3, with a distance of only 0.1 AU, whereas KP.3_F59S had a distance of 1.3 AU relative to KP.3 (**Fig. 3B, Fig. S1B**). Again, the glycosylation mutants of XEC and KP.3.1.1 clustered distinctly from their parental variants, similar to the patten observed in the bivalent cohort (**Fig. 3A-B, Fig. S1B**).

In XBB.1.5-vaccinated hamsters (**Fig. 3C, Fig. S1C**), the overall antigenic distances in this cohort were much smaller than the other cohorts, as we have shown previously (11, 14, 21). However, the JN.1-lineage variants still clustered distinctly from D614G, with an average distance of 2.8 AU (ranged 2.5-3.0 AU). XEC again exhibited one of the largest distances from D614G, i.e., 3.0 AU compared to 2.7 AU for KP.3.1.1. Differences between individual mutations were less clear due to closer clustering, though the single mutation KP.3_F59S clustered distinctly than T22N and was farther away from both KP.3 and JN.1. The glycosylation mutants were also much closer to their parental variants than in the other cohorts (**Fig. 3C, Fig. S1C**).

### XEC spike exhibits notably lower fusogenicity, which is rescued by removal of the acquired glycosylation mutation T22N

We next examined the fusogenicity of the spikes of interest. Briefly, HEK293T cells were co-transfected with the spike of interest together with GFP then co-cultured with target cells, either HEK293T-ACE2 (**Fig. 4A-B**) or CaLu-3 (**Fig. 4C-D**). As we have shown previously, Omicron spikes exhibited notably lower fusion than D614G in both cell lines (6, 11, 14, 18, 19, 21, 24). In HEK293T-ACE2 cells, XEC exhibited the lowest fusogenicity, with a 1.9-fold drop relative to JN.1 (p < 0.0001), a 0.9-fold drop relative to KP.3, and a 1.5-fold decrease relative to KP.3.1.1 (p < 0.01). This drop appeared to be driven by the T22N mutation which exhibited a 1.3-fold drop relative to KP.3 (p < 0.01), compared to the F59S, which remained comparable to KP.3 (**Fig. 4A-B**). The glycosylation mutants for XEC and KP.3.1.1 both afforded notable increases in fusion relative to their parental spikes, with XEC_S24A increasing fusion by 2.0-fold relative to XEC (p < 0.0001) and KP.3.1.1_T33A increasing fusion by 1.2-fold relative to KP.3.1.1 (p < 0.05) (**Fig. 4A-B**).

**Figure 4:**
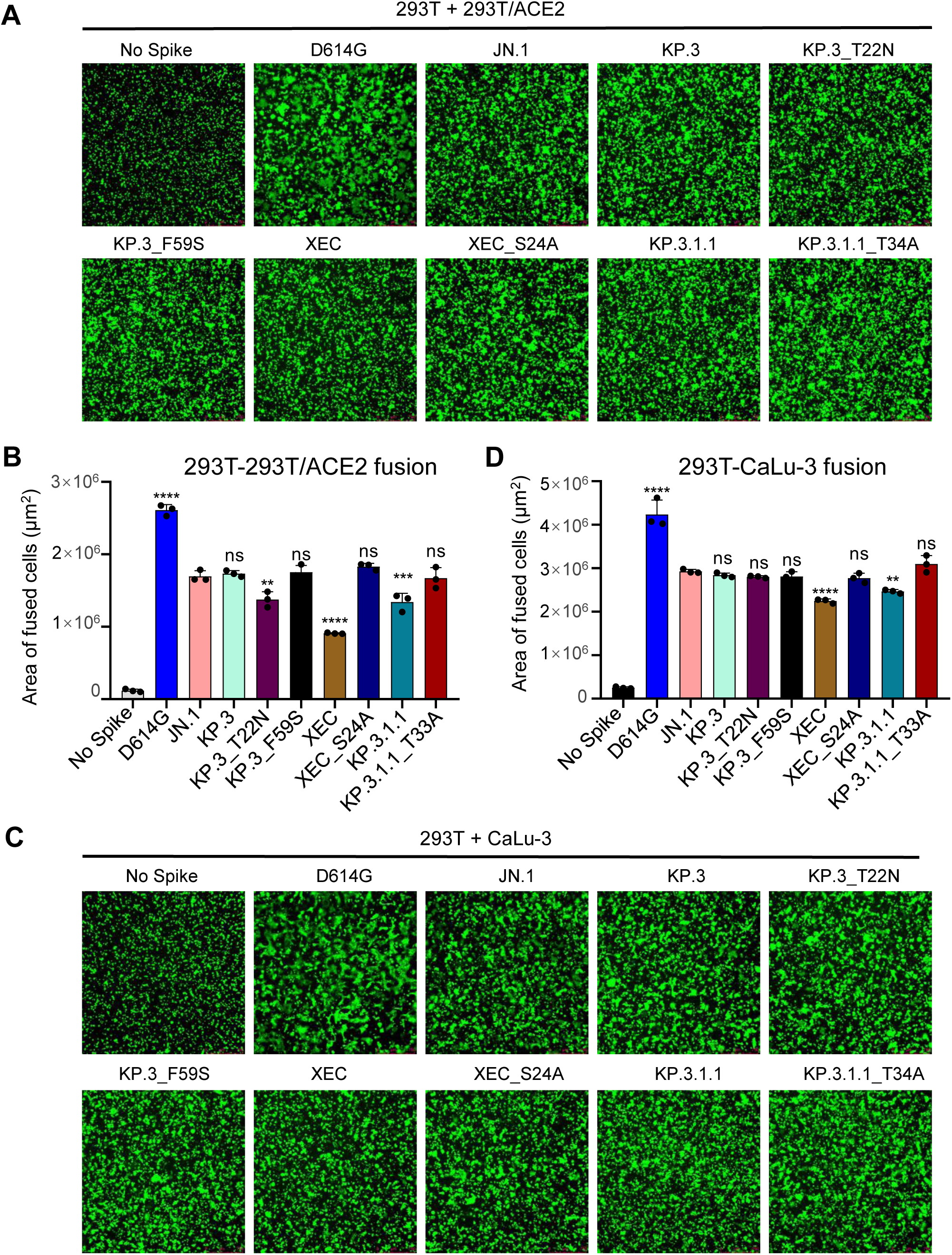
XEC exhibits decreased fusogenicity in HEK293T-ACE2 and CaLu-3 cells. Fusion of spikes was determined in HEK293T-ACE2 cells **(A-B)** and CaLu-3 cells **(C-D)**. Representative images of fusion are depicted for 293T-ACE2 **(A)** and CaLu-3 **(C)**, and quantification of total areas of fusion across 3 images are represented for in **(B)** 293T-ACE2 and **(D)** CaLu-3 cells. Areas of fused cells were determined using microscope software (see Methods). Plots represent means with standard deviation with significance determined in comparison to JN.1. Significance was calculated using repeated measures one-way ANOVA and represented as ns p > 0.05, **p < 0.01, ****p < 0.0001.

The same overall trends were observed in CaLu-3 cells (**Fig. 4C-D**). XEC exhibited the lowest cell-cell fusion, with a 1.3-fold drop relative to JN.1 (p < 0.0001). This decrease was almost comparable to KP.3.1.1, which had a drop of 1.2-fold relative to JN.1 (p < 0.01). Of note, neither T22N nor F59S drove the decreased fusion in CaLu-3 cells, both remaining comparable to JN.1 (**Fig. 4C-D**). Importantly, loss of the acquired glycosylation mutation recovered fusion activity of both XEC and KP.3.1.1, with XEC_S24A exhibiting a 1.2-fold increase relative to XEC (p < 0.01) and KP.3.1.1_T33A exhibiting an increase of 1.3-fold relative to KP.3.1.1 (**Fig. 4C-D**). Overall, similar to that in HEK293T-ACE2 cells, XEC and KP.3.1.1 exhibited decreased cell-cell fusion compared to their ancestral JN.1, and the acquisition of glycosylation at the NTD appeared to play an important role.

### Surface expression, S1 shedding, and spike processing of XEC and its derived mutants

To assess whether differences in cell-cell fusion could be attributed to differences in spike expression on the surface of cells, we conducted flow cytometry to measure levels of spike on the surface of HEK293T cells. As shown in **Fig. 5A-B**, expression levels of all spikes were comparable, with XEC, KP.3_F59S, KP.3.1.1, and KP.3_T33A exhibiting a modest increase, whereas KP.3_T22N showed a modest decrease. We also determined the spike processing in the transfected HEK293T cells by immunoblotting cell lysates and observed no differences for XEC, mutants and tested variants (**Fig. 5C**). Hence, the differences in cell-cell fusion cannot be explained by the spike expression level on the cell surface or efficiency in its processing by furin.

**Figure 5:**
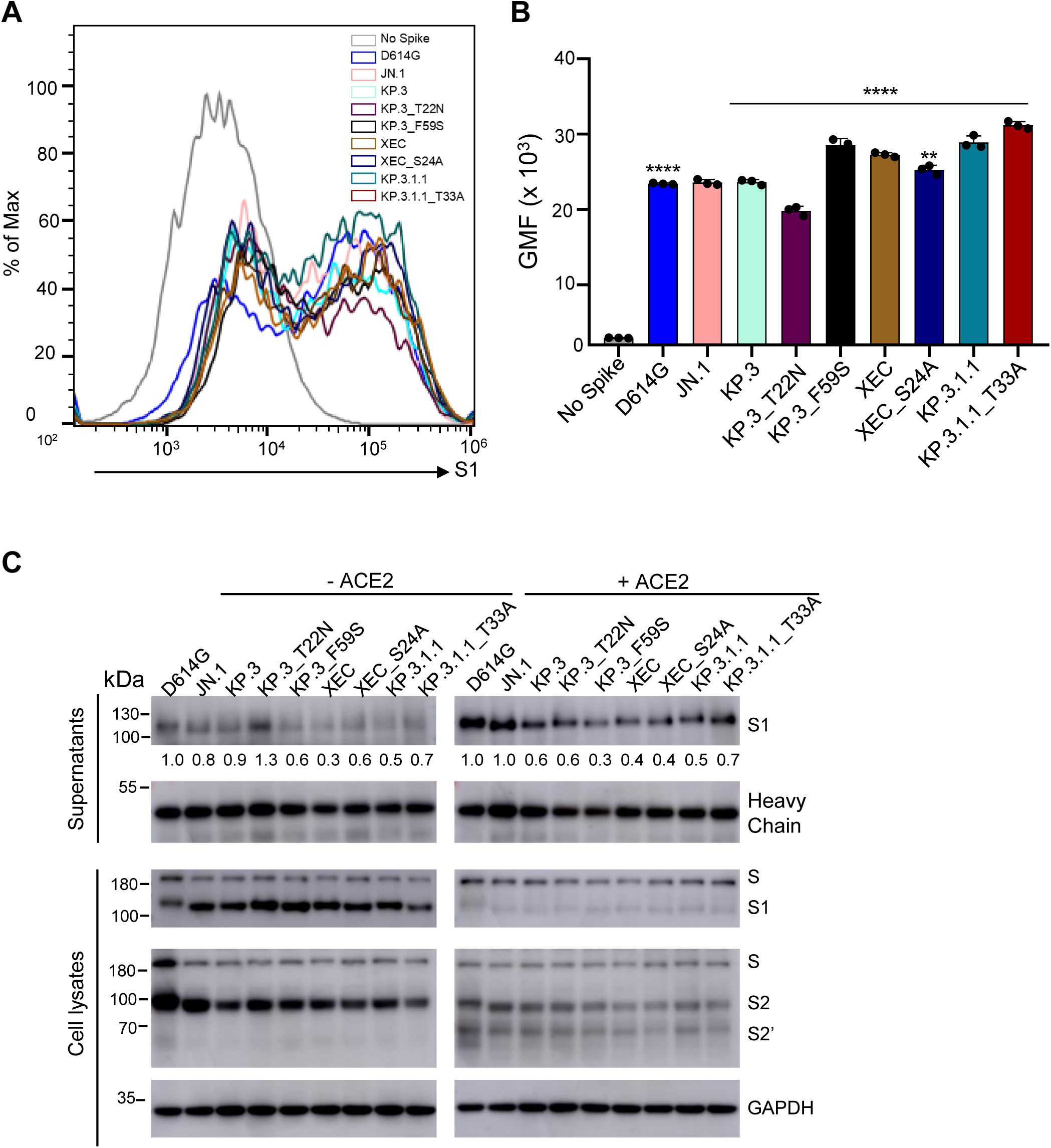
The surface expression, processing, and S1 shedding of XEC spike and related variants. **(A-B)** Surface expression of spikes on the membrane of HEK293T cells was determined using flow cytometry using anti-S1 antibody. **(A)** Representative histograms and **(B)** plots of averaged geometric mean fluorescence intensity (GMFI) are depicted. Plots represented geometric mean fluorescence intensities with standard deviation. Significance in **(B)** was determined using repeated measures one-way ANOVA and represented as ****p < 0.0001.**(C)** Spike expression in transfected cells and S1 shedding. HEK293T cells were transfected with spike constructs of interest and treated with or without sACE2 (10 μg/ml) for 4 h. Cell culture media and lysates were collected, with shed S1 proteins being immunoprecipitated with an anti-S1 antibody. Cell lysates with or without sACE2 were blotted with anti-S2, anti-S1 and anti-GAPDH antibodies, and relative signals were quantified by NIH ImageJ (30) by setting the value of JN.1 to 1.0. Images from one representative experiment are shown.

Given our recent results showing that the NTD DelS31 mutation increases spike stability potentially by acquiring a N-linked glycosylation and thus contributing to the decreased cell-cell fusion of LB.1, KP.2.3 and KP.3.1.1 variants (14), we determined S1 shedding of XEC and its derived mutants, including those ablating the glycosylation mutation, i.e, KP.3_S24A and KP.3.1.1_T33A. We transfected HEK293T cells with the spike protein of interest, and treated cells with or without 10 µg/ml soluble ACE2 (sACE2) for 4 hours. Culture media and cell lysates were collected and subjected to immunoblotting using an anti-S1 antibody. As shown previously (14), treatment of cells with sACE2 dramatically enhanced S1 shedding across all spike variants tested (**Fig. 5C**; compare the signal between the left and right panels). Notably, XEC and its KP.3_F59S showed a reduced level of S1 shedding compared to their parental KP.3, with and without the sACE2 stimulation (**Fig. 5C**). While the loss of its glycosylation mutation, i.e., XEC_S24A, slightly increased S1 shedding in the absence of sACE2, this was not the case when sACE2 was present. Interestingly, we consistently observed an increased S1 shedding for KP.3.1.1_T33A relative to KP.3.1.1, both in the presence and absence of sACE2 (**Fig. 5C**), strongly suggesting that the glycosylation mutation by DelS31 in KP.3.1.1 spike is likely responsible, at least in part, for the decreased S1 shedding. The heavy chain signals of the anti-S1 antibody used for the pulldown of the spike were comparable, indicating that differences in S1 shedding were not due to the input amount of the anti-S1 antibody used (**Fig. 5C**). Immunoblotting of the cell lysates was performed using anti-S1 and anti-S2 antibodies, respectively, revealing comparable levels of spike expression and cleavage into S1 and S2 in the transfected cells (**Fig. 5C**). As would be expected, we observed dramatically decreased S1 signals, yet increased S2’ intensity, after cells were treated with ACE2, reflecting the ACE2-mediated triggering of S1 shedding and subsequent spike activation (**Fig. 5C**; compare intensities between the left and right panels). These results together showed that XEC has decreased S1 shedding, which is likely attributed to the F59S mutation, rather than T22N; yet DelS31 mutation in KP.3.1.1 largely contributes to its stability by acquiring a N-linked glycosylation in the NTD.

### Molecular modeling of key mutations in XEC spike

Molecular modeling of key mutations in the XEC spike provides valuable insights into how these changes might contribute to immune escape, cell-cell fusion and spike stability (**Fig. 6A-C**). The T22N mutation introduces an N-linked glycosylation sequon at position 22, resulting in addition of a glycan at this site. This glycan protrudes outward, partially overlapping a critical antibody epitope within the NTD. As a result, antibodies such as C1717 (25), which typically recognize this region, may experience reduced efficacy due to the interference caused by this glycan modification (**Fig. 6B**). This modification likely facilitates viral immune escape by reducing the efficacy of neutralizing antibodies targeting the NTD, thereby promoting evasion of pre-existing immunity. The F59S mutation, on the other hand, alters the hydrophobic interactions between F59 and adjacent residues, including F32, F59, and L293 (**Fig. 6A**). This disruption may induce an allosteric conformational change in the spike protein, thus impacting its overall stability and possibly make the spike more prone to premature activation or destabilization. Additionally, the F59S mutation introduces new hydrogen bonds with N30 (**Fig. 6A**), further modifying the local structure and potentially affecting the spike’s ability to properly transition between its functional conformations. This structural change could also impair the binding of antibodies like 4-33 (26), which rely on interactions with the hydrophobic phenylalanine residue at position 59, reducing antibody affinity and contributing to immune resistance (**Fig. 6C**). Collectively, these two mutations in the XEC variant would diminish the binding of NTD-targeting antibodies, enhancing the virus’s ability to escape immune recognition and neutralization.

**Figure 6.**
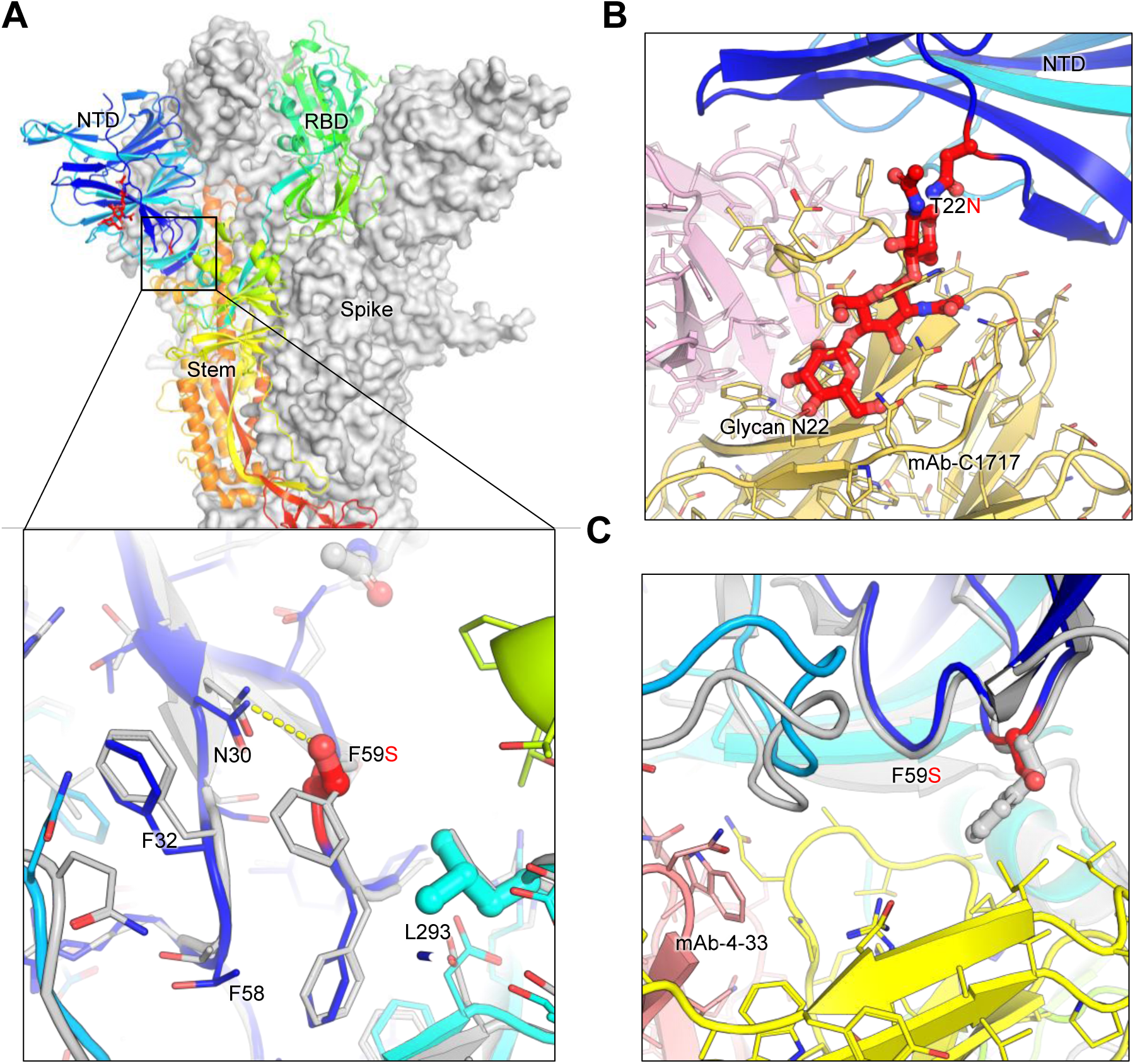
Structural modeling of key mutations in XEC spike. **(A)** Structural representation of the spike protein domains, with the location of NTD mutations T22N and F59S highlighted. The spike is shown with two protomers displayed as a grey surface and one as a ribbon in rainbow colors. Inset: The F59S mutation alters its side-chain interaction with several nearby hydrophobic residues, including F32, F58, and L293, while introducing a hydrogen bond with residue N30. **(B)** The glycosylation at N22 (shown as sticks) interferes with the recognition of certain NTD-targeting antibodies, such as C1717, potentially reducing antibody binding efficiency. **(C)** The F59S mutation disrupts the epitopes of NTD-targeting antibodies, such as 4-33, by abolishing the interaction with a hydrophobic cluster, thereby impairing antibody recognition and contributing to immune evasion.

## DISCUSSION

As new cases of COVID-19 continue to arise worldwide, it remains critical to assess emerging variants for changes in spike biology, most notably the activity of nAb escape and virus infectivity. The recombinant XEC variant is currently on the rise in the US and globally, so it is important to establish whether it will overtake the currently dominating KP.3.1.1, and to determine the underlying mechanisms for its selective advantage. Here, we show that XEC displays an increased infectivity compared to its parental KP.3, though it is still lower than KP.3.1.1 in both HEK293T-ACE2 and CaLu-3 cells (**Fig. 1**). This finding is in agreement with results of other preprints, which showed a lower infectivity for XEC relative to KP.3.1.1 in HOS-ACE2-TMPRSS2 (27) and CaLu-3 cells (13) and a comparable infectivity in Vero cells (13, 17). Importantly, we found that the single mutation F59S at the NTD largely accounts for the increased infectivity, while the T22N mutation located in the same region does not have a significant impact, a finding is corroborated by others (27). Deep mutational scanning on the XBB.1.5 spike has revealed that mutation F59S could afford a modest increase in ACE2 binding (28), though other studies have observed comparable ACE2 binding between KP.3, KP.3.1.1, and XEC (13, 17). Despite the mixed findings, these observations together highlight the importance of NTD mutations and their role in dictating critical aspects of spike biology. Crucially, we showed in lower airway epithelial CaLu-3 cells that XEC still has notably lower infectivity relative to D614G, similar to all other previous Omicron variants (6, 17–19), although the tropism and pathogenesis of XEC and KP.3.1.1 in vivo remains to be investigated.

One common feature of XEC and KP.3.1.1 is mutations in the NTD that create new glycosylation sites. The DelS31 mutation in KP.3.1.1 is predicted to generate a glycosylation site on residue N30, while the T22N mutation in XEC is expected to create a glycosylation site on residue N22. To investigate the role of glycosylation in spike biology and nAb escape, we introduced mutations that would ablate glycosylation sites, e.g. KP.3.1.1_T33A and XEC_S24A. Western blotting analysis reveals that S1 bands for the glycosylation mutants migrate slightly faster than the parental variants (**Fig. 5C**), indicating a likely loss of the spike glycosylation in these sites due to the DelS31 and T22N mutation. Interestingly, ablating the glycosylation site in KP.3.1.1 caused notably decreased infectivity (**Fig. 1**), and partially restored nAb neutralization, especially in the bivalent vaccinees and BA.2.86/JN.1 patient cohorts (**Fig. 2**), as well as shortened the antigenic distance (**Fig. 3**). These results are corroborated by Liu et al., who demonstrated that the glycosylation site in XEC can dictate inhibition by soluble ACE2 and RBD-targeting monoclonal antibodies (17). We also discovered that this loss of the glycosylation mutation in both KP.3.1.1 and XEC led to increased cell-cell fusion (**Fig. 4**), which is accompanied by enhanced S1 shedding (**Fig. 5**). Together, these results indicate that the NTD modification by glycosylation critically dictates the spike stability, virus infectivity, and nAb neutralization.

Although the NTD does not directly interact with the ACE2 receptor, it plays an essential role in maintaining the spike protein’s conformation and dynamics. Our homology modeling (**Fig. 6**) suggests that the mutations in the NTD of XEC spike, T22N and F59S, may impact the spike stability and viral infectivity, though the precise mechanisms require further experimental validation. The T22N mutation introduces an N-linked glycosylation site at position 22, potentially hindering antibody recognition and promoting immune evasion. Experimental data indicate that this glycosylation reduces cell-cell fusion activity, likely due to a steric hindrance from the added glycan, which could restrict the conformational flexibility necessary for efficient fusion. However, this effect is modest and can vary with cellular context or other unexamined factors. Conversely, the F59S mutation disrupts hydrophobic interactions between F59 and adjacent residues, introducing conformational changes that increase spike flexibility and enhancing its adaptability for receptor binding. Additionally, F59S could influence allosteric regulation of the receptor-binding domain (RBD) ↓ as the NTD and RBD are structurally linked, and mutations in the NTD can affect the RBD’s tendency to adopt the “up” conformation essential for ACE2 binding. Overall, increased local flexibility from F59S would encourage the RBD to remain in this open state, enhancing ACE2 binding and therefore viral infectivity and nAb escape. We must emphasize that the full impact of T22N and F59S on antibody recognition, cell-cell fusion, and infectivity remains speculative and awaits experimental validation. Nonetheless, understanding the complex interplay between these structural alterations and functional outcomes related to the spike NTD is critical and can provide valuable insights into virus-host interaction and vaccine design.

Overall, our study highlights the importance of studying emerging variants of SARS-CoV-2, particularly as evolution shifts to less characterized regions of spike, especially the NTD. We have shown the critical role of NTD mutations in dictating aspects of spike biology that can in turn impact vaccine efficacy and disease manifestation. Importantly, recent data by Arora et al. demonstrated that JN.1 booster vaccination can induce nAb titers against KP.3.1.1 and XEC, but they are still lower than titers against JN.1 (13), suggesting new formulations will still have to be considered moving forward.

## LIMITATION OF STUDY

In our study, we made use of lentivirus pseudotyped vectors bearing the spike protein of interest. Ideally, these assessments would be made using authentic SARS-CoV-2 variants. However, we have previously validated this pseudotyped virus system alongside infectious virus (29), and believe the timeliness of this work justifies their use. Additionally, our sera cohorts are relatively limited in size because of regulatory constraints. We have previously applied similarly sized cohorts and obtained reliable results that are valuable to inform the government regulatory agency for updating COVID-19 vaccines. Despite this, the limited sample size can have influences on assessment of significance.

## ACKNOWLEDGEMENTS

We thank the Clinical Research Center and Center for Clinical Research Management of The Ohio State University Wexner Medical Center and The Ohio State University College of Medicine in Columbus, Ohio, especially Breona Edwards, Evan Long, J. Brandon Massengill, Francesca Madiai, Dina McGowan, and Trina Wemlinger, for collecting and processing the samples. We also thank Sarah Karow, Gabrielle Swoope, Daniela Farkas, and the Critical Care Clinical Trials team of The Ohio State University for sample collection and other supports. We acknowledge Ashish R. Panchal, Mirela Anghelina, Soledad Fernandez, and Patrick Stevens for their assistance in providing the sample information of the first responders and their household contacts. We thank Peng Ru and Lauren Masters for sequencing and Xiaokang Pan for bioinformatic analysis. S.-L.L., D. J., R.J.G., L.J.S. and E.M.O. are supported by the National Cancer Institute of the NIH under award no. U54CA260582. The content is solely the responsibility of the authors and does not necessarily represent the official views of the National Institutes of Health. This work was also supported by a fund provided by an anonymous private donor to OSU. K.X. was supported by NIH grants U01 AI173348 and UH2 AI171611. J.N.F. was supported by a Glenn Barber Fellowship from the Ohio State University College of Veterinary Medicine. M.C. was supported by an NIH T32 training grant (T32AI165391) and Dean’s Graduate Enrichment Fellowship at The Ohio State University. J. L. was supported by NIH R01AI090060 and P01AI175399. J.S.B was supported by NIH K08 HL169725. R.J.G. was additionally supported by the Robert J. Anthony Fund for Cardiovascular Research and the JB Cardiovascular Research Fund, and L.J.S. was partially supported by NIH R01 HD095881.

## AUTHOR CONTRIBUTIONS

S.-L.L. conceived and directed the project. R.J.G led the clinical study/experimental design and implementation. P.L. performed the experiments and data processing and analyses. Y.X. and K.X. performed molecular modeling and data analyses. Y.L. assisted experiments. C.C.H, M.C., and J.L. provided hamster serum samples and associated information. D.J. led SARS-CoV-2 variant genotyping and DNA sequencing analyses. C.C., J.S.B., J.C.H., R.M., and R.J.G. provided clinical samples and related information. P.L., J.N.F. and S.-L.L. wrote the paper. Y.-M.Z, L.J.S., E.M.O. provided insightful discussion and revision of the manuscript.

## FIGURE LEGENDS

**Table S1: Details of neutralization cohorts.** Demographic information and vaccine details are listed for each neutralization cohort.

**Table S2: Antigenic distance units to variants of interest relative to D614G or JN.1 (Related to** Fig. 3). Antigenic distance (AD) values were determined using Microsoft PowerPoint for each of the variants relative to D614G or JN.1 and listed for each cohort.

## METHODS

### Lead contact

Dr. Shan-Lu Liu can be reach at with any questions or requests for reagents.

### Materials Availability

Materials can be requested from the lead contact.

### Data and Code Availability

This study reports no original code. Raw data can be requested from the lead contact.

## EXPERIMENTAL MODEL AND SUBJECT DETAILS

### Vaccinated and patient cohorts

Full demographic information and details of vaccination can be found in **Table S1.**

This study makes use of 3 cohorts, the first of which were bivalent vaccinated healthcare workers at the OSU Wexner Medical center (n=8). These individuals received three doses of monovalent WT spike mRNA vaccine and one dose of bivalent (WT+BA.4/5) spikes mRNA vaccine. Four received Moderna and 4 Pfizer. 7 individuals received a third dose of vaccine (4 Moderna, 3 Pfizer) while 1 individual did not receive a third dose. Five individuals were administered the Pfizer formulation of the bivalent vaccine while 5 received the Moderna formulation. Blood was collected between 23-97 days post bivalent dose administration. Individuals ranged from 27-46 years old with a median of 39, 5 males and 3 females were recruited. Samples were collected under IRB protocols 2020H0228, 2020H0527, and 2017H0292.

The next cohort were patients at the OSU Wexner Medical Center that were either admitted to the ICU during the BA.2.86/JN.1 wave of infection in Columbus, OH (11/23/2024-8/11/2024) (n=5) or collected from first responders and household contacts in the STOP-COVID cohort that were symptomatic during that time period (n=4). RT-PCR was used to confirm COVID-19 positivity and infecting variant was determined using next gen sequencing (Artic v5.3.2, IDT, Coralville, IA and Aritc v4.1 primers, Illumina, San Diego, CA). Ages ranged from 34-77 with a median of 51. 4 females and 5 males were recruited to this cohort. Samples were collected under IRB protocols 2020H0527, 2020H0531, 2020H0240, and 2020H0175.

The final cohort were golden Syrian hamsters that were vaccinated with two doses of recombinant Mumps vaccine expressing the XBB.1.5 spike (n=9). The vaccine was administered intranasally at 1.5 x 10^5^ PFU twice three weeks apart. All hamsters were 15 weeks old and had blood collected 2 weeks after the booster dose. Studies were conducted under IRB protocols 2009A1060-R4 and 2020A00000053-R1.

### Cell lines and maintenance

Cells used in this study included HEK293T (ATCC, RRID: CVCL_1926), HEK293T-ACE2 (BEI Resources, RRID: CVCL_A7UK), and CaLu-3 cells (ATCC, Cat #30-2003). HEK293T cells were maintained in DMEM (Sigma Aldrich, Cat #11965-092) with 10% fetal bovine serum (Thermo Fisher, Cat #F1051) and 0.5% penicillin/streptomycin (HyClone, Cat #SV30010). CaLu-3 cells were maintained in EMEM (ATCC, Cat #30-2003) supplemented the same way. To passage, cells were first washed with PBS then detached using 0.05% trypsin + 0.53 mM EDTA (Corning, Cat #27106). Cells were incubated at 37°C with 5.0% CO_2_.

## METHOD DETAILS

### Plasmids

All spike plasmids are in the pcDNA3.1 backbone and are tagged at the C-terminal end with single FLAG tags. D614G was synthesized via restriction enzyme cloning at KpnI and BamHI by GenScript Biotech. The JN.1 spike was made in-house through site-directed mutagenesis of BA.2.86 (synthesized by GenScript). KP.3, KP.3.1.1, XEC, and individual mutants were all generated in-house through site-directed mutagenesis of corresponding parental variants. All constructs were confirmed by DNA sequencing. pNL4-3_inGluc is an HIV-1 lentiviral vector used for pseudotyping (31).

### Lentiviral pseudotype production and infectivity measurement

Lenviral pseudotypes were produced through via polyethyleneimine transfection (Transporter 5 Transfection Reagent, Polyscienes, Cat #26008-5) of 293T cells. Cells were transfected in a 2:1 ratio of vector to spike and supernatant containing vectors was collected 48 and 72 hours post-transfection. This supernatant was clarified and used to infect HEK293T-ACE2 or CaLu-3 cells. CaLu-3 were subjected to spin-inoculation at 1,650 x g for 1hr to enhance attachment. To assess relative infection, media containing secreted *Gaussia* luciferase was collected off target cells and combined with an equal volume of *Gaussia* luciferase subtstrate (0.1 M Tris pH 7.4, 0.3 M sodium ascorbate, 10 µM coelenterazine). The readout was collected on a BioTek Cytation 5 Imaging Reader. Readings were collected 48 and 72 hours post infection.

### Lentiviral pseudotype neutralization assay

Vectors produced above were used a neutralization assay as described previously (29). Briefly, vectors were diluted to normalize infectivity then incubated with serially diluted sera (1:80, 1:320, 1:1,280, 1:5,120, 1:20,480). This mixture was then used to infect HEK293T-ACE2 cells. Readouts were collected as described above and used to determine NT_50_ values through least squares nonlinear regression against a no sera control via GraphPad v10 (San Diego, CA).

### Antigenic cartography map generation

Antigenic mapping was carried out via the Racmacs v1.1.35 program. The corresponding GitHub (https://github.com/acorg/Racmacs/tree/master) was used to run the raw NT_50_ values through the program using R (Vienna, Austria). The program performed log2 transformation on the values then plotted them in a distance table. This table was then used to perform multidimensional scaling and plot the individual spike antigens as circles and individual sera samples as squares in two-dimensional space where 1 antigenic distance unit (AU) represents about a 2-fold change in average NT_50_ between antigens. Racmacs optimizations were kept on default and maps were exported using the “view(map)” function. Maps were labeled and AU between antigens determined using Microsoft Office PowerPoint.

### Cell-cell fusion

HEK293T cells were transfected with spike of interest alongside GFP then co-cultured with the target HEK293T-ACE2 or CaLu-3 cells. Cells were co-cultured together for either 6.5 hours (HEK293T-ACE2) or 4 hours (CaLu-3) then imaged using a Leica DMi8 fluorescence microscope. To determine the areas of fused cells, the Leica X Applications Suite was used to outline areas of fusion based on GFP signal and calculate the space with each area. Scale bars in images represent 150 µM. Three representative images were taken for each variant and used for quantification, one image was then chosen for presentation in Fig. 4.

### Spike surface expression and processing

HEK293T cells were transfected spike of interest and used for staining with a polyclonal S1 antibody (Sino Biological, T62-40591, RRID:AB_2893171) followed by anti-Rabbit-IgG-FITC secondary (Sigma, F9887, RRID:AB_259816) to determine difference in spike expression on the cell membrane. Flow cytometry was performed using an Attune NxT flow cytometer and data was analyzed using FlowJo v10.8.1.

Lysates from spike transfected HEK293T cells were collected using RIPA buffer (Sigma Aldrich, R0278) supplemented with protease inhibitor (Sigma, P8340) and subjected to SDS-PAGE on a 10% polyacrylamide gel, followed by transfer onto a PVDF membrane. Immunoblotting was performed with polyclonal S1 antibody (Sino Bio, T62-40591, RRID:AB_2893171), polyclonal S2 antibody (Sino Biological, T62-40590, RRID:AB_2857932), and anti-GAPDH (Proteintech, 10028230). Spike antibodies were probed with anti-rabbit-IgG-HRP (Sigma, Cat#A9169, RRID:AB_258434), anti-GAPDH antibodies were probed with anti-mouse-IgG-HRP (Sigma, Cat#A5728, RRID:AB_258232). HRP chemiluminescence was read out using Immobilon Crescendo Western HRP substrate (Millipore, WBLUR0500) on a GE Amersham Imager 600.

### S1 shedding

HEK293T cells were transfected with spike of interest. Following 24 hours of transfection, cells were then treated with or without 10 μg/mL soluble ACE2 (Sino Biological, Cat# 10108-H08H-B) and incubated for 4 hours at 37°C to induce S1 shedding. Cell culture media and lysates were then collected. A pulldown was performed on the media using 10 μL of protein A/G-conjugated anti-S1 beads (Santa Cruz, sc-2003) overnight to obtain shed S1, which was detected by immunoblotting using anti-S1 antibody (Sino Bio, T62-40591, RRID:AB_2893171) alongside cell lysate samples.

### Structural modeling and analysis

Structural modeling was conducted using the SWISS-MODEL server (32). Glycosylation modifications at residue N22 were incorporated using the Coot program to simulate potential structural alterations. Published cryo-EM structures (PDB: 8D55, 8D5A, 8WLY, 9FJK, 8Y5J, 7UAR, 8CSJ, 8OYT) served as templates for this analysis. The effects of key mutations, including T22N and F59S, on spike protein interactions, stability, and immune evasion were assessed. The resulting models were visualized and analyzed using PyMOL to investigate how these mutations may influence the spike’s functional properties and ability to escape immune recognition.

### Statistical analyses

Statistical analyses in this work were conducted using GraphPad Prism 10. NT_50_ values were calculated by least-squares fit non-linear regression. Error bars in Figures 1D, 1E, 4B, 4D and 5B represent means ± standard errors. Error bars in Figures 2A, 2C, and 2E represent geometric means with 95% confidence intervals. Statistical significance was analyzed using log10 transformed NT_50_ values to better approximate normality (Figures 2A, 2C and 2E), and multiple groups comparisons were made using a one-way ANOVA with Bonferroni post-test. Cell-cell fusion was quantified using the Leica X Applications Suite software (Figures 4A and 4C). S processing was quantified by NIH ImageJ (Figure 5).

