## Supplemental Table for "Immune Evasion, Cell-Cell Fusion, and Spike Stability of the SARS-CoV-2 XEC Variant: Role of Glycosylation Mutations at the N-terminal Domain"

### Table S1. Bivalent-vaccinated HCW and BA.2.86/JN.1-wave first responder cohorts

| | Bivalent Health Care Workers (n=8) | BA.2.86/JN.1 Wave Patients (n=9) |
|---|---|---|
| **Age in Years at Sample Collection [Median (Range)]** | 39 (27-46) | 51 (34-77) |
| **Gender** [n (% of Total)] | | |
| Male | 5 (63%) | 5 (56%) |
| Female | 3 (37%) | 4 (44%) |
| **Sample Collection Window** | Dec. 2022 | Nov. 2023-Aug.2024 |
| **Vaccine status** [n (% of Total)] | NA | |
| 1-dose Pfizer | NA | 1 (10%) |
| 2-dose Moderna | NA | 2 (20%) |
| 4-dose Moderna | NA | 1 (10%) |
| 1-dose Moderna +1-dose Pfizer bivalent | NA | 1 (10%) |
| 1-dose Pfizer +1-dose Pfizer bivalent | NA | 2(20%) |
| 2-dose Pfizer +1-dose Pfizer bivalent | 1 (12.5%) | NA |
| 3-dose Pfizer +1-dose Moderna bivalent | NA | 1 (14.3%) |
| 3-dose Pfizer +1-dose Pfizer bivalent | 3 (37.5%) | NA |
| 3-dose Moderna +1-dose Moderna bivalent | 4 (50%) | 1 (14.3%) |
| Days from last vaccination | NA | 621 (34-1033) |
| Days post the bivalent dose for recipients | 65 (23-97) | NA |
| **COVID-19 positive** [n (% of Total)] | 8 (80%) | 9 (100%) |
| Days before sample collection [(Median Range)] | 324 (182-994) | 7 (1-10) |
| **Infected Variants** | | |
| JN.1/BA.2.86 | NA | 2 (22%) |
| Undetermined | NA | 8 (78%) |

Summary of the demographic information for two cohorts used for neutralization experiments depicted in Figure 2. “NA” means the category is not applicable to the cohort.

| Bivalent HCWs | | | |
|---|---|---|---|
| AD (D614G) | | AD (JN.1) | |
| JN.1 | 4.9 | D614G | 4.9 |
| KP.3 | 5.3 | KP.3 | 1.0 |
| KP.3_T22N | 5.4 | KP.3_T22N | 1.3 |
| KP.3_F59S | 6.8 | KP.3_F59S | 3.6 |
| XEC | 7.5 | XEC | 3.3 |
| XEC_S24A | 6.7 | XEC_S24A | 4.8 |
| KP.3.1.1 | 7.3 | KP.3.1.1 | 5.0 |
| KP.3.1.1_T33A | 6.1 | KP.3.1.1_T33A | 1.2 |

| BA.2.86/JN.1-wave patients | | | |
|---|---|---|---|
| AD (D614G) | | AD (JN.1) | |
| JN.1 | 3.4 | D614G | 3.4 |
| KP.3 | 4.1 | KP.3 | 1.7 |
| KP.3_T22N | 4.3 | KP.3_T22N | 1.7 |
| KP.3_F59S | 4.9 | KP.3_F59S | 3.1 |
| XEC | 5.7 | XEC | 3.4 |
| XEC_S24A | 5.2 | XEC_S24A | 4.2 |
| KP.3.1.1 | 5.3 | KP.3.1.1 | 2.9 |
| KP.3.1.1_T33A | 4.9 | KP.3.1.1_T33A | 1.8 |

| XBB.1.5-monovalent hamsters | | | |
|---|---|---|---|
| AD (D614G) | | AD (JN.1) | |
| JN.1 | 2.7 | D614G | 2.7 |
| KP.3 | 2.7 | KP.3 | 0.2 |
| KP.3_T22N | 2.8 | KP.3_T22N | 0.2 |
| KP.3_F59S | 2.5 | KP.3_F59S | 0.7 |
| XEC | 3.0 | XEC | 0.7 |
| XEC_S24A | 3.0 | XEC_S24A | 0.5 |
| KP.3.1.1 | 2.7 | KP.3.1.1 | 0.6 |
| KP.3.1.1_T33A | 2.8 | KP.3.1.1_T33A | 0.7 |

**Figure S1**
